# Identifying novel regulators of placental development using time series transcriptomic data and network analyses

**DOI:** 10.1101/2022.05.17.492330

**Authors:** Ha T. H. Vu, Haninder Kaur, Kelby R. Kies, Rebekah R. Starks, Geetu Tuteja

## Abstract

The placenta serves as a connection between the mother and the fetus during pregnancy, and provides the fetus with oxygen, nutrients, and growth hormones. However, the regulatory mechanisms and dynamic gene interaction networks underlying early placental development are understudied. Here, we generated RNA sequencing (RNA-seq) data from mouse fetal placenta tissues at embryonic day (e) 7.5, e8.5 and e9.5 to identify genes with timepoint-specific expression, then inferred gene interaction networks to analyze highly connected network modules. We determined that timepoint-specific gene network modules associated with distinct developmental processes, and with similar expression profiles to specific human placental cell populations. From each module, we obtained hub genes and their direct neighboring genes, which were predicted to govern placental functions. We confirmed that four novel candidate regulators identified through our analyses regulate cell migration in the HTR-8/SVneo cell line. Upon conclusion of this study, we were able to predict several novel regulators of placental development using network analysis of bulk RNA-seq data. Our findings and analysis approaches will be valuable for future studies investigating the transcriptional landscape of early placental development.

## Introduction

The placenta is a transient organ that has critical roles during pregnancy, such as the transportation of oxygen and nutrients to the fetus, waste elimination, and the secretion of growth hormones. Placental defects are associated with devastating complications including preeclampsia and fetal growth restriction, which can lead to maternal or fetal mortality [1], [2]. Therefore, it is fundamental to understand the mechanisms of placental development.

Due to ethical considerations as well as the opportunity for genetic manipulation, mouse models are frequently used when investigating early placental development. Like humans, mice have a hemochorial placenta [3], meaning that maternal blood directly comes in contact with the chorion. Although there are certain differences between the mouse and human placenta [3], [4], they do share common regulators and signaling pathways involved in placental development [4], [5]. For example, Ascl2/ASCL2 [6], [7] and Tfap2c/TFAP2C [8] are required for the trophoblast (TB) cell lineage in both mouse and human models. Additionally, the HIF signaling pathway is conserved as it regulates TB differentiation in both mouse and human systems [9].

Mouse placental development begins around embryonic day (e) 3.5 when the trophectoderm (TE) layer forms [5]. The TE differentiates into different TB populations at e4.5, which eventually leads to the formation of the ectoplacental cone (EPC) [10]. Between e7.5 and e9.5, the establishment of blood flow to the fetus begins, resulting in highly dynamic changes in placental cell composition. At e7.5, the EPC is comprised of TB cells [3], organized into the inner and peripheral populations, with the inner cells actively proliferating and differentiating, while the outer cells can be invasive and interact with the decidua [10]. Around e8.5, chorioallantoic attachment occurs, during which the chorion layer joins with the allantois [11]. As a result, the e8.5 mouse fetal placenta includes cells from the EPC, chorion and allantois [12]. From e9.5 onwards, the mouse fetal placenta is composed of distinct layers, the trophoblast giant cell (TGC) layer, the junctional zone (spongiotrophoblast and glycogen TB cells), and the labyrinth zone (chorion TB cells, syncytiotrophoblast I and II cells, fetal endothelium, and spiral artery TGCs) [13], [14]. Within the labyrinth layer, there is a dense network of vasculature where nutrients and oxygen are transported and exchanged. Several individual regulators of the processes active between e7.5 and e9.5 have been identified, as reviewed in [5], [15]–[18].

In addition to identifying individual regulators governing placental functions, it is also important to determine how these regulators potentially interact with other genes as networks. In order to identify novel regulators or infer gene interactions underlying developmental processes, unbiased whole genome transcriptomic data can be used. Previous studies that utilized transcriptomics in the developing mouse placenta were either focused on analysis of one timepoint, or focused on analysis of multiple -omics data [19]–[22]. Other studies of gene expression in human placenta across trimesters did not infer full gene interaction networks and instead focused on transcription factors [23], [24]. Single-cell analysis has been used to investigate cell-type specific gene expression in the placenta; however, these studies do not predict regulators underlying specific placental development processes [25], [26].

Here, we generated RNA sequencing (RNA-seq) data from mouse fetal placental tissues at e7.5, e8.5 and e9.5. We then carried out clustering, differential expression, and network analyses to infer gene interactions and predict novel regulators of placental development. We further demonstrated that our network constructions could be used to infer cell populations in the mouse placenta at the three timepoints. Finally, we conducted in vitro validation experiments and confirmed that several genes we identified have a role in regulating TB cell migration.

## Results

### 1. Genes associated with distinct placental processes show timepoint-specific expression

We generated and analyzed transcriptomic data from fetal placental tissues at e7.5, e8.5 and e9.5 to identify genes regulating distinct processes during placental development. Based on the stages of placental development and the cell types present at each stage, we predicted that genes with highest expression at e7.5 would be involved in TB proliferation or differentiation; genes with highest expression at e8.5 would have a role in chorioallantoic attachment; and genes with highest expression at e9.5 would have a role in the establishment of nutrient transport. Indeed, we observed that previously identified regulators of TB proliferation and differentiation (e.g., Ascl2 [6], [27], Gjb5 [28]), chorioallantoic attachment (e.g., Ccnf [29], Itga4 [30]), and nutrient transport (e.g., Gjb2 [31], Igf2 [32]) showed timepoint-specific patterns that matched with our predictions (Figure 1A). Next, we performed hierarchical clustering (see Materials and Methods) to determine if protein-coding transcripts would cluster into groups that displayed timepoint-specific expression. From this analysis, we obtained three groups of transcripts in which the median expression was highest at e7.5 (8242 transcripts, equivalent to 5566 genes), e8.5 (8091 transcripts, equivalent to 5536 genes) and e9.5 (7238 transcripts, equivalent to 5347 genes) (Figure 1B, Supplementary Table S1). Hereafter, these groups are referred to as hierarchical clusters.

**Figure 1:**
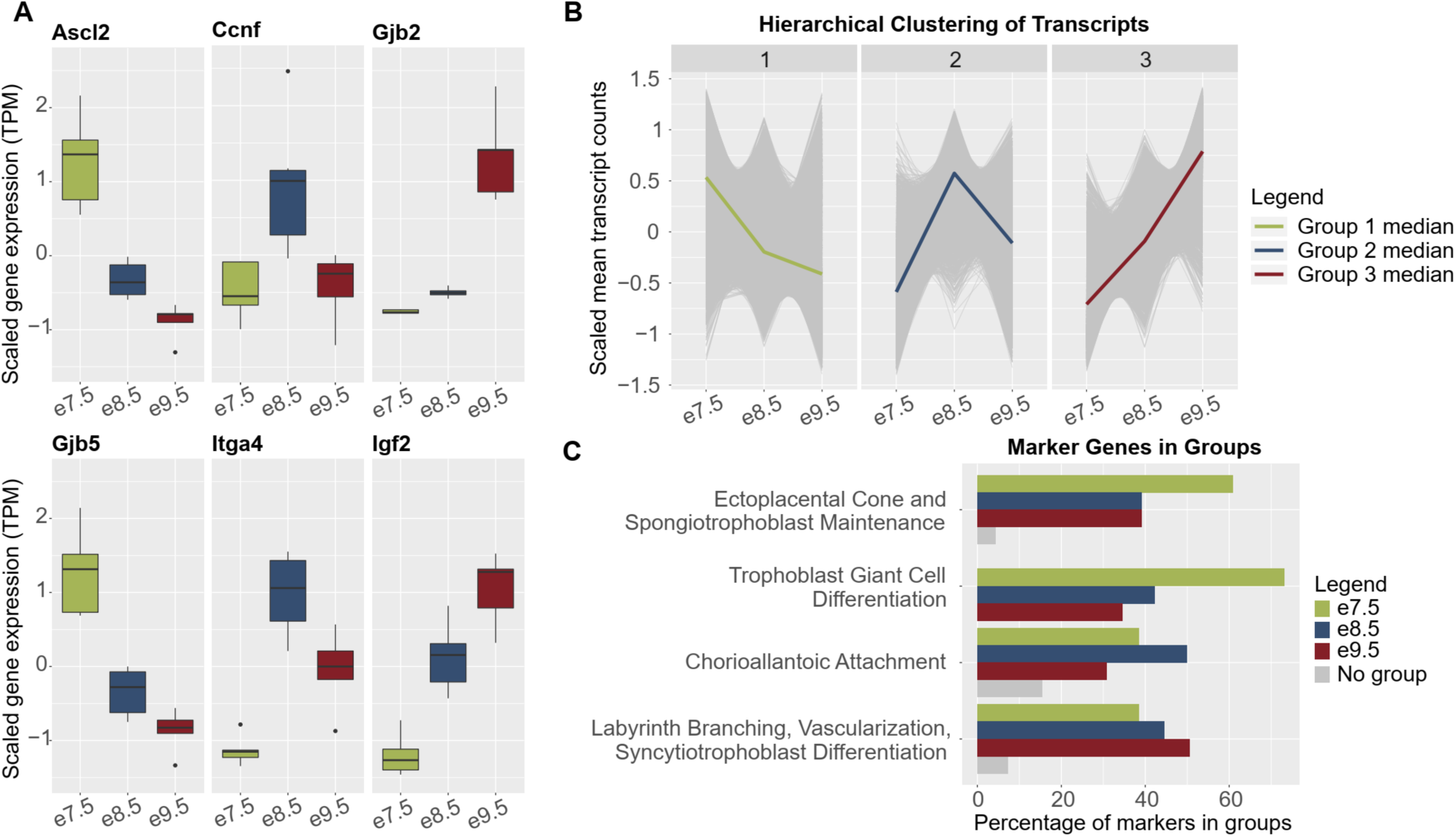
Genes associated with distinct placental processes show timepoint-specific expression. A. Boxplots of scaled mean expression (in transcripts per million, TPM) of marker genes showing timepoint-specific patterns. Ascl2 and Gjb5, expected to peak at e7.5, markers of trophoblast proliferation and differentiation [6], [34]; Ccnf and Itga4, expected to peak at e8.5, markers of chorioallantoic attachment [29], [30]; Gjb2 and Igf2, expected to peak at e9.5, markers of nutrient transport [31], [32]. B. Line charts of scaled mean raw counts of transcripts in hierarchical clusters showing group median expression levels peak at each timepoint. C. Bar plots showing that timepoint-associated hierarchical clusters captured most genes underlying distinct placental processes. Markers of timepoint-associated placental processes were obtained from previously published review articles [5], [15]–[18]. Green, markers in hierarchical cluster with median expression level highest at e7.5; blue, markers in hierarchical cluster with median expression level highest at e8.5; dark red, markers in hierarchical cluster with median expression level highest at e9.5; grey, markers in no hierarchical clusters.

To evaluate the computational robustness and biological significance of the hierarchical clusters, we carried out additional analyses. First, we validated the grouping patterns using three different algorithms, K-means clustering, self-organizing maps, and spectral clustering, and observed similar trends (Supplementary Figure S1). Second, we determined how the genes in each cluster relate to processes of placental development. From previously published review articles [5], [15]–[18], we acquired gene sets associated with distinct processes, namely ectoplacental cone and/or spongiotrophoblast maintenance (expected to be most active at e7.5, when the EPC is still in a highly proliferative state [10]), trophoblast giant cell differentiation (expected to be more active at e7.5 because the mouse placenta at e8.5 and e9.5 includes more differentiated TB subtypes [3], [12]–[14]), chorioallantoic attachment (expected to be most active at e8.5 [11]), and labyrinth branching, vascularization and syncytiotrophoblast differentiation (expected to be most active at e9.5, after these processes have initiated [18]) (Supplementary Table S2). Indeed, we observed that the e7.5 hierarchical cluster captured the most genes in the ectoplacental cone and spongiotrophoblast maintenance and trophoblast giant cell differentiation group; the e8.5 hierarchical cluster included the most genes in the chorioallantoic attachment group; and the e9.5 hierarchical cluster included the most genes in the labyrinth branching, vascularization and syncytiotrophoblast differentiation group (Figure 1C, Supplementary Table S2). Together, these data demonstrate that hierarchical clustering can be used to obtain transcript groups that are associated with relevant biological processes at each timepoint, but is not sufficient to fully distinguish processes that may have varied activity levels throughout time.

To this end, and because hierarchical clustering is sensitive to small perturbations in the datasets [33], we carried out differential expression analysis (DEA) and identified transcripts and genes with the strongest changes over time (Supplementary Figure S2, Supplementary Table S3). After combining results from hierarchical clustering and DEA, we defined timepoint-specific gene groups (see Materials and Methods for gene group definitions, Supplementary Figure S2) and obtained 922 e7.5-specific genes, 915 e8.5-specific genes, and 1952 e9.5-specific genes (Supplementary Table S4). These timepoint-specific groups were further analyzed to predict gene interactions and functions.

### 2. Network analysis reveals potential regulators of developmental processes in the placenta

To predict interactions amongst timepoint-specific genes and subset timepoint-specific genes into regulatory modules, we used the STRING database [35] and GENIE3 [36] (see Materials and Methods). With the two approaches of network inference, we were able to predict networks of genes by means of previously published experimental results and text-mining of available publications (STRING), as well as de novo computational analysis with random forest-based methods (GENIE3). We then carried out network sub-clustering with the GLay algorithm [37] (see Materials and Methods) and identified four network modules at e7.5, six at e8.5, and eight at e9.5 (Supplementary Table S5). To determine if the networks were associated with distinct processes of placental development, we used gene ontology (GO) enrichment analysis.

Compared to e8.5 and e9.5 networks, e7.5 networks had a higher rank or fold change for the GO terms “inflammatory response” and “female pregnancy” (Figure 2A). The term “morphogenesis of a branching structure”, which can be expected following choriollantoic attachment around e8.5, was not enriched at e7.5, but was enriched in multiple e8.5 and e9.5 networks (e8.5_1_STRING, e8.5_2_GENIE3, e9.5_1_STRING, e9.5_1_GENIE3, e9.5_2_STRING, and e9.5_2_GENIE3). After chorioallantoic attachment finishes, nutrient transport is being established. Accordingly, we observed the following enrichments: “endothelial cell proliferation” (highest ranked in e9.5_2_STRING), “lipid biosynthetic process” (only significant after e7.5, highest ranked in e9.5_3_STRING), “cholesterol metabolic process” (only significant after e7.5, highest ranked in e9.5_2_GENIE3 and e9.5_3_STRING), and “response to insulin” (only significant after e7.5, highest ranked in e9.5_1_GENIE3). “Tube morphogenesis”, “vasculature development” and cell migration related terms (“positive regulation of cell migration” and “epithelium migration”) are observed in networks at all timepoints, although these terms are not consistently ranked in all networks. Supplementary Table S6 includes the full list of significant terms observed.

**Figure 2:**
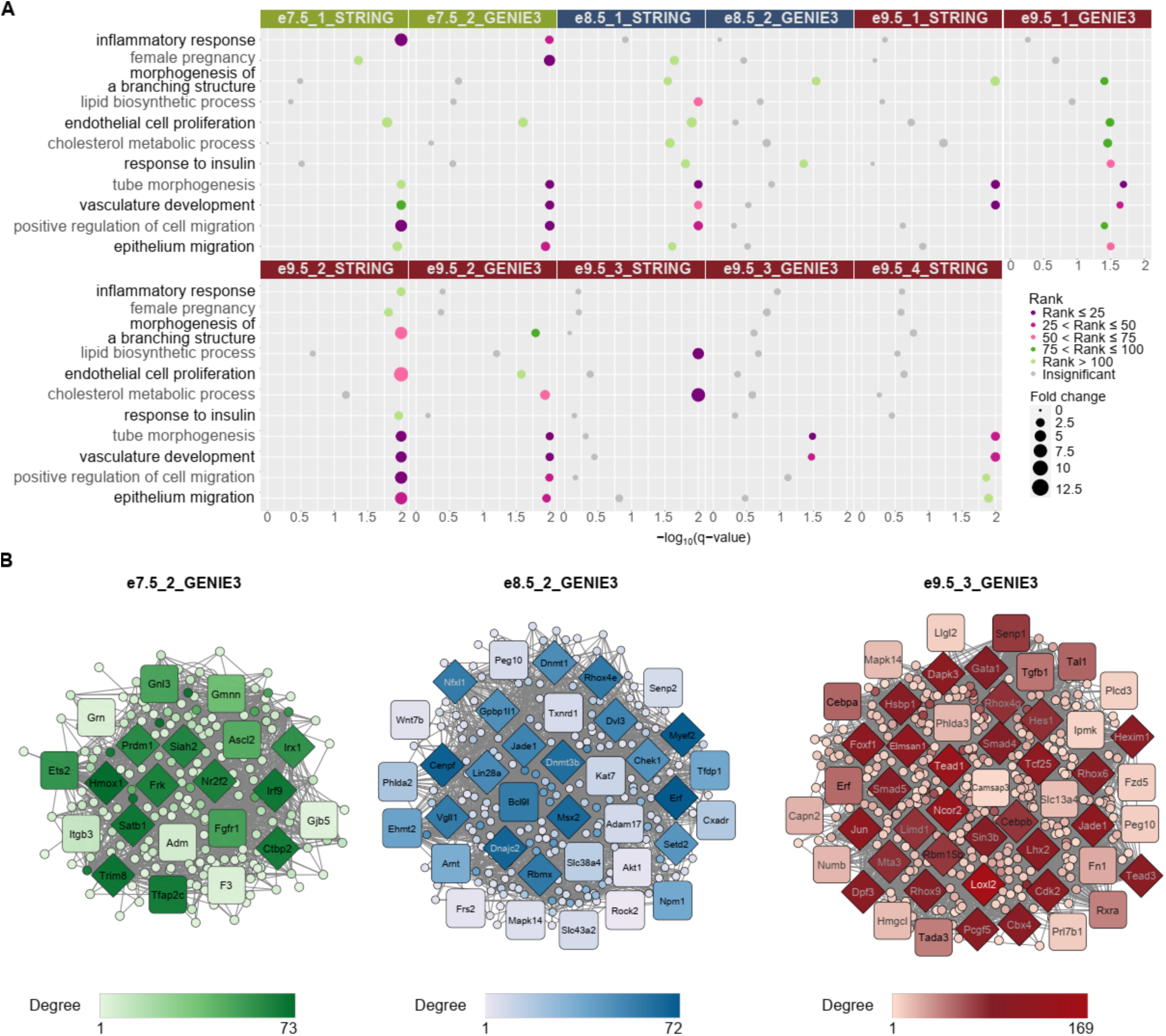
Network analysis identifies gene modules with relevant functions and reveals potential regulators of placental development. A. Gene ontology (GO) analysis of networks demonstrates the association of gene sets with placental development processes. Only selected terms are shown. Dot colors correspond to ranks of the terms in each analysis; dot sizes correspond to fold change. A GO term is considered enriched if its q-value ≤ 0.05, fold change ≥ 2, and the number of observed genes ≥ 5. For full GO enrichment analysis, see Supplementary Table S6. B. Network analysis highlights potential regulators of placental development. Only a subset of networks with enriched terms from (A) are shown. Square shaped nodes: genes annotated with placental functions on the MGI database (see Materials and Methods). Diamond shaped nodes: hub genes. Color: the darker the color is, the higher the node’s degree centrality is.

We predicted that hub genes, defined to be nodes with high degree, closeness, and shortest path betweenness centrality in the networks (see Materials and Methods), could be potential regulators of developmental processes in the placenta. We first determined if the hub genes from each network described in the previous paragraph were annotated with placental functions using the Mouse Genome Informatics (MGI) database [38] (see Materials and Methods, Table 1, Supplementary Table S7). In the network e7.5_1_STRING, we identified seven hub genes (Table 1), none of which were annotated to terms related to placental development according to the MGI database. In the network e7.5_2_GENIE3 (Figure 2B), ten hub genes were identified, three of which were annotated in the MGI database, and are required for TB proliferation, differentiation, migration or invasion, namely Nr2f2 [39], Prdm1 [40], and Ctbp2 [41] (Table 1, Supplementary Table S7). However, upon further literature search, we found that the two networks’ hub genes include genes that have an established role in placental development that are not annotated by MGI, such as Mmp9 (e7.5_1_STRING), Hmox1 and Satb1 (e7.5_2_GENIE3), which are required for proper implantation, TB differentiation and invasion [42], [43], [44]. Other hub genes could be novel regulators of placental functions. One example is Frk, a hub gene of the e7.5_2_GENIE3 network, which had been suggested to inhibit cell migration and invasion in human glioma [45] and retinal carcinoma cells [46], but has not been studied in early placental development.

**Table 1:**
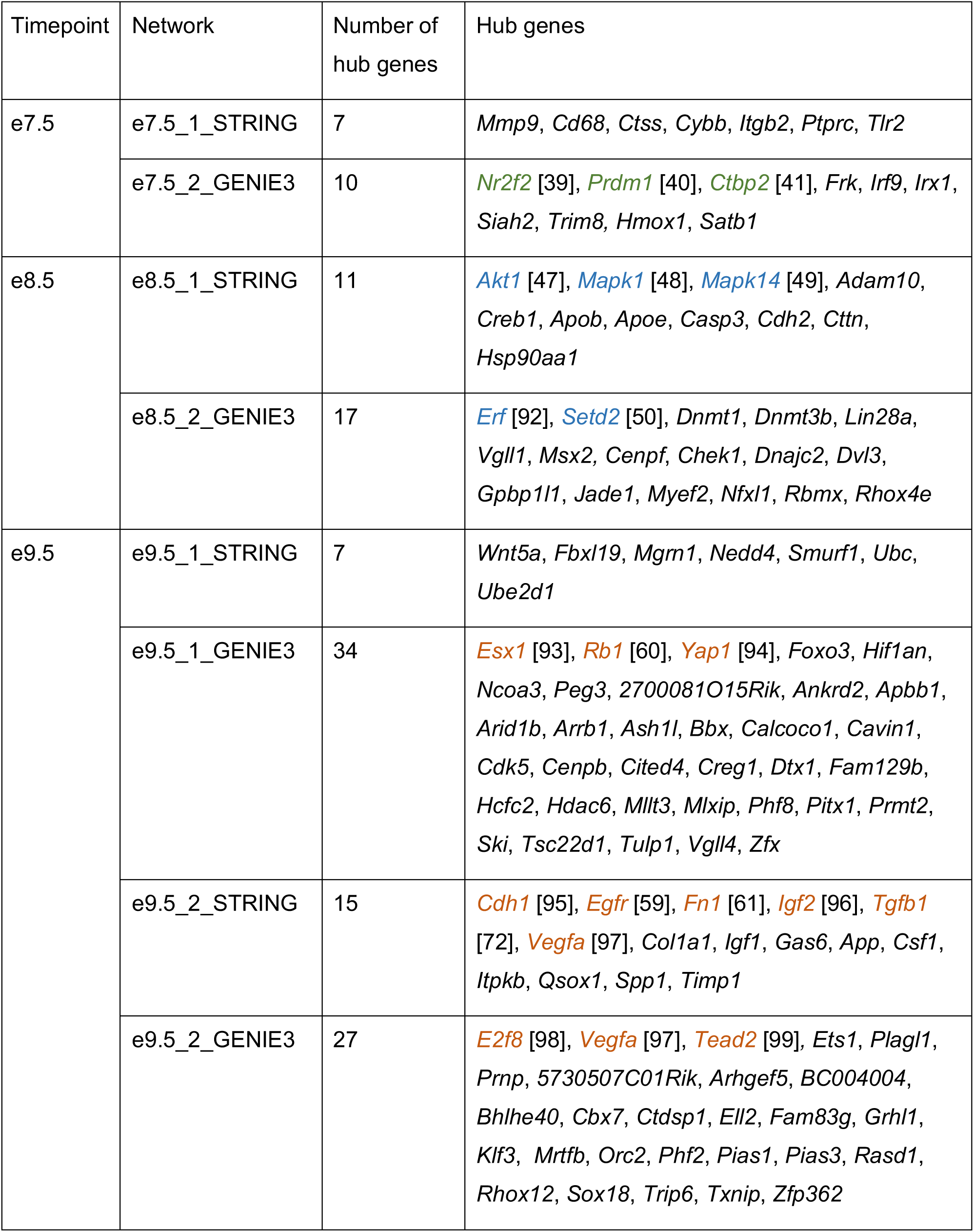
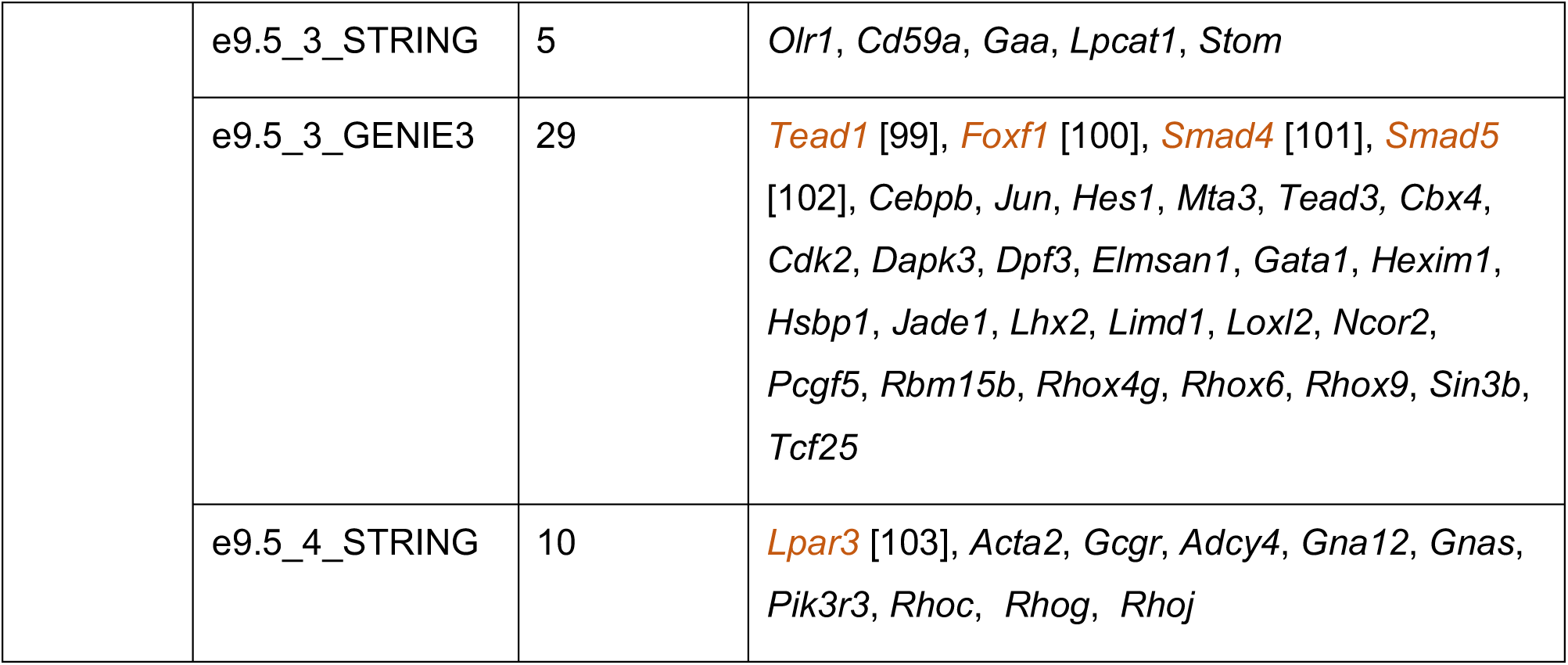
Hub genes associated with each network. Colored genes are ones that have annotated roles in placental development (see Materials and Methods); green, e7.5-specific genes; blue, e8.5-specific genes; brown, e9.5-specific genes.

At e8.5, hub genes included both novel and known genes in placental development or chorioallantoic attachment. For example, in the network e8.5_1_STRING, 11 hub genes were identified, of which three were associated with placental development according to the MGI database. These genes, Akt1, Mapk1, and Mapk14, all have a role in placental vascularization [47]–[49] (Table 1, Supplementary Table S7). For the network e8.5_2_GENIE3 (Figure 2B), there were 17 hub genes identified (Table 1, Supplementary Table S7), with three genes annotated with placenta related terms in MGI and associated with placental development processes such as placental vascularization (Setd2 [50]). Both networks’ hub genes include genes that are not annotated to placental function related terms, but have been studied in the context of placental development such as Adam10 [51] and Creb1 [52] (e8.5_1_STRING), Dnmt1 [53], Dnmt3b [54], Lin28a [55], Vgll1 [56], and Msx2 [57] (e8.5_2_GENIE3). An example of a novel gene is Jade1 (hub node of e8.5_2_GENIE3), which has been found to have high expression in extraembryonic ectoderm and TB cells and hence may play roles in placental vascularization by interacting with VHL [58], but has not been tested functionally in placental tissues.

From the e9.5 networks, we identified 127 hub genes of which 16 have been annotated as having a role in placental development in the MGI database (Table 1, Supplementary Table S7). For instance, in e9.5_1_GENIE3, e9.5_2_STRING, and e9.5_4_STRING, hub genes that regulate labyrinth layer development include Egfr [59] and Rb1 [60], and hub genes that regulate placental vasculature development include Fn1 [61] and Vefga [62]. The hub genes from these networks again include genes that are not annotated with placental development terms, but have a known role in placental development such as Wnt5a [63] (e9.5_1_STRING), Ets1 [64] (e9.5_2_GENIE3), and Cebpb [65] (e9.5_3_GENIE3). There are also hub genes known to be important for placental nutrient transport such as Igf2 [32], and other genes that could be novel regulators. For example, Lhx2 is part of the mTOR signaling pathway in osteosarcoma [66], but has yet to be studied in placenta although the mTOR signaling pathway is known to be involved in nutrient transport in the placenta [67].

### 3. Timepoint-specific genes can be associated with cell-specific expression profiles of human placenta

To determine if timepoint-specific genes could capture different placental cell populations, we carried out deconvolution analysis with LinSeed [68] and inferred the cell type profiles. Briefly, LinSeed takes advantage of the mutual linearity relationships between cell-specific genes and their corresponding cells to infer the topological structures underlying cell populations of tissues. This approach would enable us to use bulk RNA-seq data to predict proportions of cell types in the mouse placenta without prior knowledge of cell type markers or matching single-cell datasets. As input to LinSeed, we used the 5000 most highly expressed genes across all timepoints (expression in TPM), from which 1413 genes were found to be statistically significant for the inference models and thus used to conduct the deconvolution analysis (see Materials and Methods, Supplementary Figure S3). As a result, we observed five cell groups which captured 99% of the variance in the placenta tissue samples (Supplementary Figure S3). Amongst these groups, e7.5 samples had the highest proportion of cell group 3, e8.5 samples had highest proportion of cell group 2, and e9.5 samples had highest proportion of cell group 5 (Figure 3A – left panel, Supplementary Table S8). Cell group 1 and cell group 4 did not have consistent cell proportions across biological replicates of a single timepoint. The identification of these cell groups could have resulted from noise introduced by both biological and technical variation, which is challenging to overcome when using a small sample size in the deconvolution analysis. Therefore, we focused on cell groups 3, 2 and 5. We identified 100 markers (see Materials and Methods) for cell group 3, 100 markers for cell group 2, and 41 markers for cell group 5. Interestingly, 95 of the 100 markers of cell group 3 are e7.5-specific genes, 45 out of 100 markers of cell group 2 are e8.5-specific genes, and 40 in the 41 markers of cell group 5 are e9.5-specific genes (Figure 3A – right panel, Supplementary Table S8). This indicates that the independent timepoint-specific gene analysis we performed in Section 2 could represent gene profiles of distinct cell populations.

**Figure 3:**
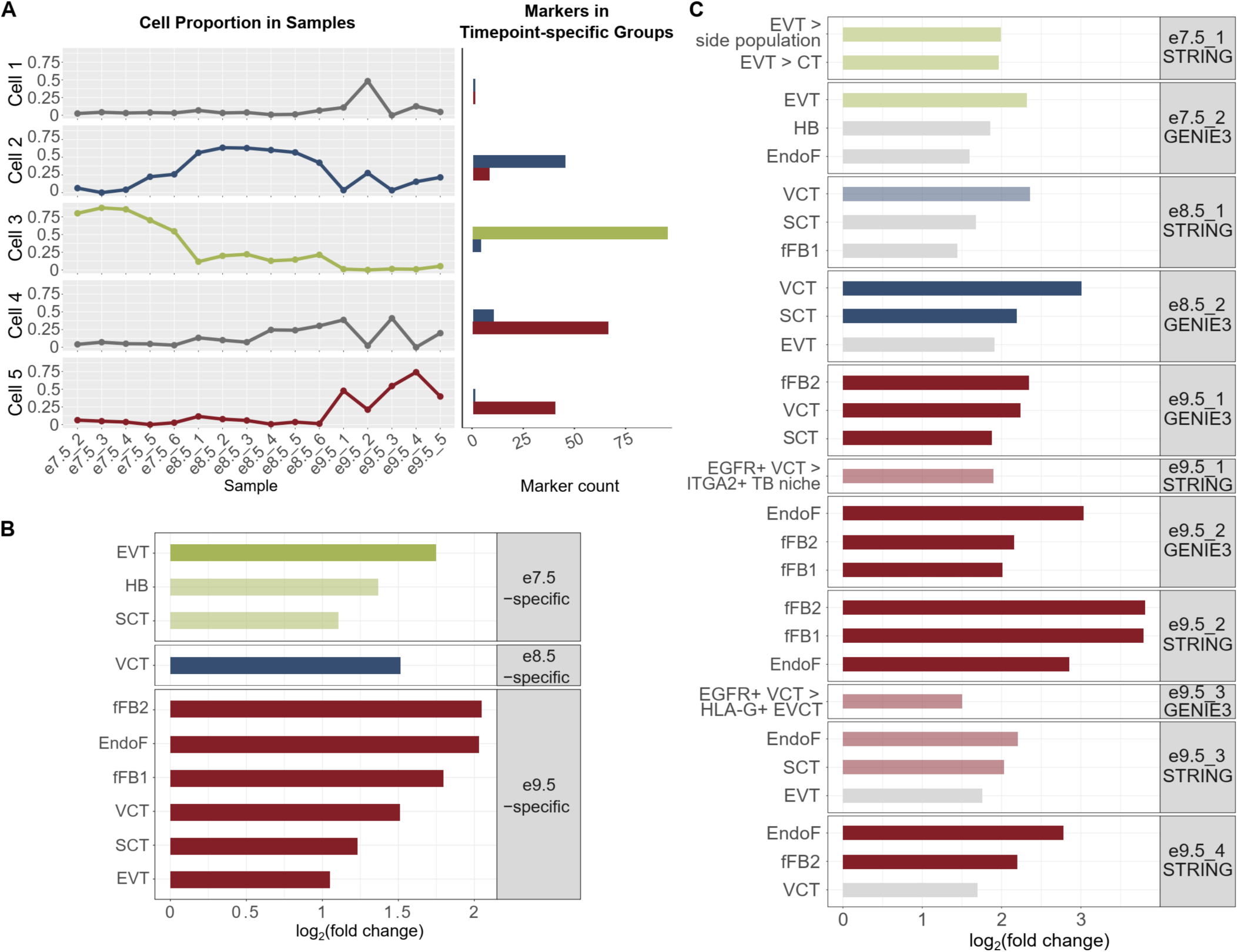
Timepoint-specific gene groups can be associated with human placenta cell-specific expression profiles. A. Deconvolution analysis using LinSeed showed five cell groups, three of which had highest proportions in e7.5 samples (group 3), e8.5 samples (group 2) and e9.5 samples (group 5). Also using LinSeed, we identified markers of each cell group and observed a high number of genes in common with timepoint-specific genes (cell group 3 with e7.5-specific genes, cell group 2 with e8.5-specific genes, cell group 5 with e9.5-specific genes). Left panel: line charts showing cell proportions in each sample; right panel: bar plots showing the number of cell markers in each timepoint-specific gene group. B. Bar plots showing that timepoint-specific genes share similar profiles to these of human placental cell populations. Enrichment analysis was carried out with PlacentaCellEnrich using 1^st^ trimester human placenta single-cell RNA-seq data to determine gene groups with cell-type specific expression. A significant enrichment has adj. p-value ≤ 0.05, fold change ≥ 2, and number of observed genes ≥ 5. The lightness of the colors corresponds to adj. p-value; the lighter colors, 0.005 < adj. p-value ≤ 0.05; the darker colors, adj. p-value ≤ 0.005. Only enrichments for cells of fetal origin are shown. Full enrichment results (including both maternal and fetal cells) are shown in Supplementary Figure S4. C. Bar plots showing that network genes share similar profiles of specific human placental cell populations. Enrichment analysis was carried out with PlacentaCellEnrich as in (B) and Placental Ontology. Grey, adj. p-value > 0.05; the lighter colors, adj. p-value ≤ 0.05; the darker colors: adj. p-value ≤ 0.005. For PlacentaCellEnrich, three fetal cell types with the lowest adj. p-values are shown. For Placenta Ontology, selected enrichments are shown. Full enrichment results (including both maternal and fetal cells and for every network) of PlacentaCellEnrich are shown in Supplementary Figure S4. Full enrichment results (for every network) of Placenta Ontology are in Supplementary Table S9. Abbreviations: SCT, syncytiotrophoblast, HB, Hofbauer cells, EVT, extravillous trophoblast, VCT, villous cytotrophoblast, EndoF, fetal endothelium, fFB1, fetal fibroblast cluster 1, fFB2, fetal fibroblast cluster 2, EVT > side population, GSE57834_extravillous_trophoblast_UP_side_population (genes upregulated in EVT compared to side population – original data from GSE57834), EVT > CT, GSE57834_extravillous_trophoblast_UP_cytotrophoblast (genes upregulated in EVT compared to cytotrophoblast – original data from GSE57834), EGFR+ VCT > ITGA2+ TB niche, GSE106852_EGFR+_UP_ITGA2+ (genes upregulated in EGFR+ villous cytotrophoblast compared to ITGA2+ proliferative trophoblast niche, original data from GSE106852), EGFR+ VCT > HLA-G+ EVCT, GSE80996_EGFR+_villous_cytotrophoblast_UP_HLA_G+_proximal_column_extravillous_cytotrophoblast (genes upregulated in EGFR+ villous cytotrophoblast compared to HLA-G+ proximal column extravillous cytotrophoblast, original data from GSE80996).

To this end, we used the PlacentaCellEnrich webtool to annotate timepoint-specific genes with human placental cell types [69]. At all timepoints, we observed enrichment suggesting the presence of TB cells. Specifically, the e7.5-specific genes were most significantly enriched for genes with extravillous trophoblast (EVT)-specific expression (log_2_(fold) = 1.75, -log_10_(adj. p-value) = 4.18), but also had enrichment for syncytiotrophoblast (SCT) (log_2_(fold) = 1.1, -log_10_(adj. p-value) = 2.09); the e8.5-specific group was only enriched for genes that had villous cytotrophoblast (VCT)-specific expression (log_2_(fold) = 1.51, -log_10_(adj. p-value) = 2.36), and the e9.5-specific group had the highest enrichment for genes with fetal fibroblast-specific expression (log_2_(fold) = 2.04, -log_10_(adj. p-value) = 22.04) (Figure 3B, Supplementary Figure S4). We note that the e9.5-specific group had enrichment for genes with cell-type specific expression in multiple cells, including endothelial cells (log_2_(fold) = 2.02, -log_10_(adj. p-value) = 18.66), VCT (log_2_(fold) = 1.5, -log_10_(adj. p-value) = 7.38), SCT (log_2_(fold) = 1.23, -log_10_(adj. p-value) = 6.93), and EVT (log_2_(fold) = 1.05, -log_10_(adj. p-value) = 3.05) (Figure 3B, Supplementary Figure S4). Together, this demonstrates that our analysis is picking up on the diverse cell populations present at e9.5 compared to e7.5.

Motivated by the fact that cell-specific expression profiles for multiple human placental cell types are enriched at e7.5 and e9.5, we hypothesized that the gene network modules at each timepoint could capture specific cell populations. Indeed, PlacentaCellEnrich analysis on e7.5_2_GENIE3 network genes was significantly enriched for genes with EVT-specific expression (log_2_(fold) = 2.32, -log_10_(adj. p-value) = 1.67) (Figure 3C, Supplementary Figure S4), but no longer with genes that have SCT-specific expression. e8.5_1_STRING and e8.5_2_GENIE3 were both enriched for genes with VCT-specific expression (log_2_(fold) = 2.35 and 3, -log_10_(adj. p-value) = 1.43 and 5.41, respectively). In addition to VCT-specific expression, e8.5_2_GENIE3 had enrichment for genes that had SCT-specific expression (log_2_(fold) = 2.19, -log_10_(adj. p-value) = 2.93) (Figure 3C, Supplementary Figure S4). At e9.5, genes in the networks e9.5_1_GENIE3 and e9.5_3_STRING showed strong enrichment for TB-specific expression, such as in SCT and VCT. On the other hand, e9.5_2_GENIE3, e9.5_2_STRING, e9.5_3_GENIE3 and e9.5_4_STRING had strong enrichment for fetal fibroblast and endothelium expression profiles (Figure 3C, Supplementary Figure S4).

For genes in network e7.5_1_STRING, e9.5_1_STRING, and e9.5_3_GENIE3, we did not observe any enrichment for fetal placental cells, possibly because not all genes in the networks are annotated in the 1^st^ trimester dataset [70] used when calculating cell enrichments in PlacentaCellEnrich. Therefore, we also used Placenta Ontology [71], which carries out enrichment tests based on different datasets than those used in PlacentaCellEnrich. With e7.5_1_STRING, in agreement with previous analyses on e7.5-specific genes or genes in e7.5_2_GENIE3 network, we observed annotations related to EVT cells being enriched, such as “EVT > side population” (log_2_(fold) = 1.99 and false discovery rate (FDR) = 0.027), and “EVT > CT” (log_2_(fold) = 1.96, FDR = 0.028) (Supplementary Table S9). With e9.5_1_STRING, the term “EGFR+ VCT > ITGA2+ TB niche” was enriched (log_2_(fold) = 1.89, FDR = 0.023), meaning there are a significant number of genes in this network that were upregulated in EGFR+ VCT compared to the ITGA2+ proliferative TB niche in 1^st^ trimester placenta. Similarly, with e9.5_3_GENIE3, we found the term “EGFR+ VCT > HLA-G+ EVCT” enriched (log_2_(fold) = 1.5, FDR = 0.043), which means there is a significant number of genes in this network that were upregulated in EGFR+ VCT compared to HGL-A+ proximal column extravillous cytotrophoblast in 1^st^ trimester placenta. In the other networks, Placental Ontology enrichment results generally agreed with PlacentaCellEnrich (Supplementary Table S9). Together, the PlacentaCellEnrich and Placenta Ontology analyses provide evidence that network analysis can be used to identify genes more likely associated with specific placental cell types.

### 4. Gene knockdown provides further evidence for a role of network genes in the placenta

As described in Section 2, we identified hub nodes, and as a result also obtained genes directly connected to the hub nodes (Supplementary Table S7). Many of the genes (23 genes at e7.5, 208 genes at e9.5) had drastic expression changes over time (having at least one transcript with fold change ≥ 5 between e7.5 and e9.5) (Supplementary Table S10), which may be more likely to have regulatory roles specific to processes or cell types associated to each timepoint. However, there were a number of hub genes and genes directly connected to the hub nodes that were differentially expressed but had lower fold changes and showed high expression across all timepoints. We predict these highly expressed genes to be generally important for TB function and processes such as cell migration, a term that was associated with multiple timepoint specific networks (Figure 2A).

To investigate this further, we performed gene knockdown and migration assays for four candidate genes from four different networks in the HTR-8/SVneo cell line, an established model for studying TB migration [72]–[74]. From the lists of hub genes and their directly connected nodes (Supplementary table S7), we obtained genes that met the following criteria: having expression levels > 5 TPM in the mouse placenta transcriptome data we generated, having expression levels > 5 FPKM (fragments per kilobase of transcript per million of mapped reads) in human TB cell lines [75] and having expression levels > 20 TPM in HTR-8/SVneo cell line [20] (Supplementary Table S7). From this list, we selected four genes: Mtdh and Siah2 (from the e7.5_1_STRING and e7.5_2_GENIE3 network, respectively), Hnrnpk (from the e8.5_2_GENIE3), and Ncor2 (from the e9.5_2_GENIE3), none of which have been studied in TB migration function. Of note, all of the networks from which we selected candidate genes were annotated to represent TB subtype populations (see Section 3).

For each of the four genes we transfected two different siRNAs, and all eight siRNAs resulted in high knockdown efficiencies (74 – 93%, Figure 4A). Next, we performed cell migration assays and observed a reduction in cell migration capacity for all four genes, as determined through quantification of integrated cell densities (Figure 4B, C, Supplementary Figure S5, Supplementary Table S11). For Siah2 and Hnrnpk, integrated densities of cells were significantly decreased upon knockdown with both siRNAs (p-value ≤ 0.05). Specifically, for Siah2, the densities reduced by 98.57% ± 0.42% (mean ± standard error) and 83.87% ± 12.1% with siRNA #1 and siRNA #2, respectively. For Hnrnpk, the densities reduced by 99.55% ± 0.09% with siRNA #1 and 98.68% ± 0.2% for siRNA #2. For Mtdh and Ncor2, the reductions were significant for one siRNA (Mtdh, siRNA #2, 98.55% ± 0.86%; Ncor2, siRNA #1, 98.11% ± 0.09%), and were fair for the other siRNA, likely due to the variable results between biological replicates (Mtdh, siRNA #1, 55.28% ± 17.22%; Ncor2, siRNA #2, 81.27% ± 14.04%). Overall, these results confirm that network analysis and gene filtering based on defined criteria can identify genes important for TB function.

**Figure 4:**
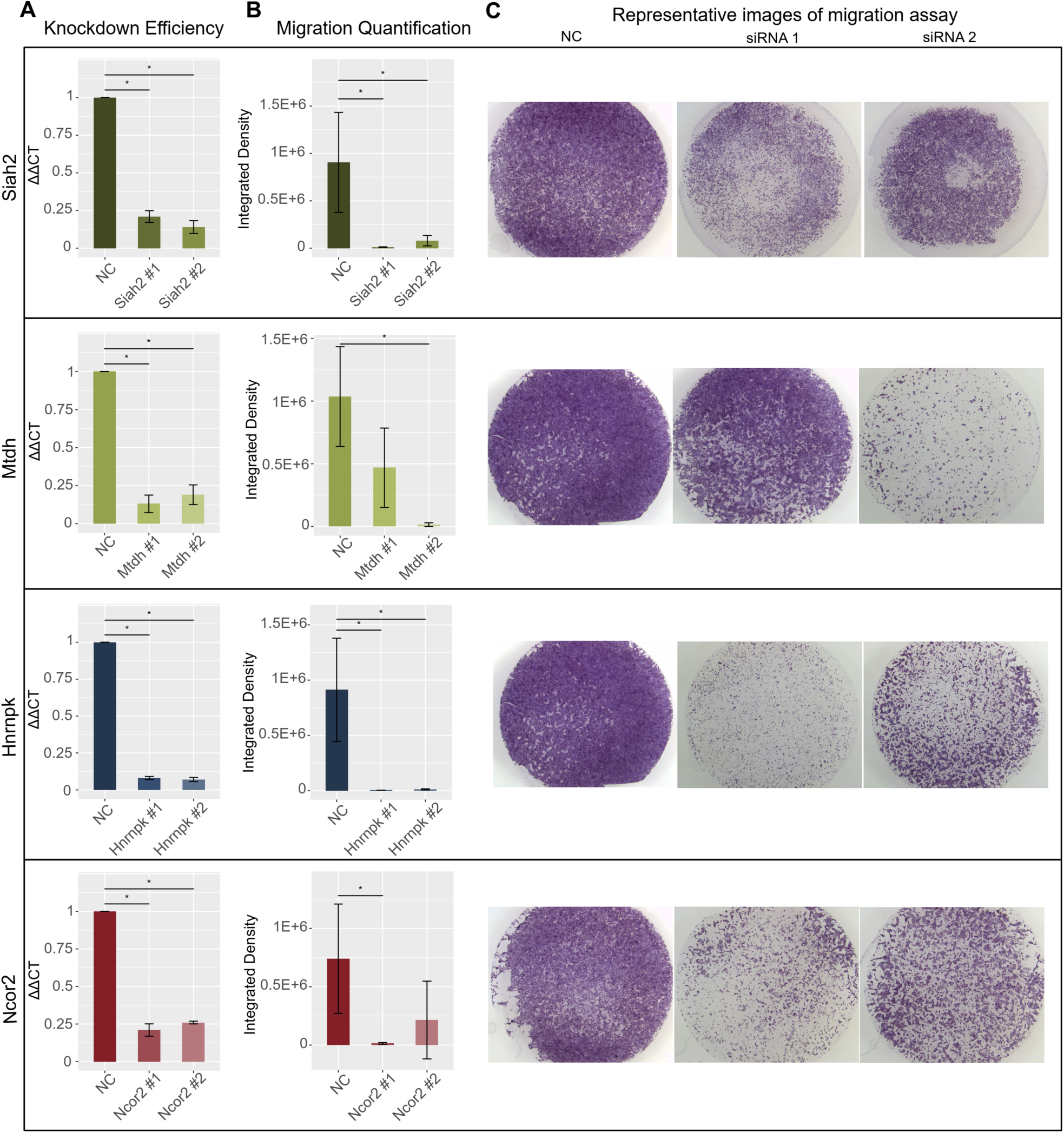
Gene knockdown of selected network genes showing reduction in cell migration capacity. Panels correspond to four genes, Siah2, Mtdh, Hnrnpk and Ncor2. Each condition, negative control (NC), siRNA #1 and siRNA #2, had three biological replicates. Error bars show standard deviation. A. Bar plots showing that gene expression was significantly reduced after knockdown (KD) compared to NC. GAPDH was used for normalization of all four genes’ expression (ΔCT). Percent KD were calculated with the ΔΔCT method. Values shown were normalized to the NC siRNAs. Y-axis shows ΔΔCT value. Details of KD efficiencies, siRNAs, and primer sequences can be found in Supplementary Table S11 and S12. (*) indicates p-value < 0.05. B. Bar plots showing significant reduction in the integrated density of cells after knockdown (KD) compared to NC samples. Y-axis shows integrated densities of cells in NC samples, samples KD with siRNA #1 of each gene, and samples KD with siRNA #2 of each gene. Details of integrated densities can be found in Supplementary Table S11. (*) indicates p-value < 0.05. C. Representative images of migration assays. Left, NC samples; middle, siRNA #1 samples; right, siRNA #2 samples.

## Discussion

Placenta development involves multiple processes that are active during different stages of gestation. Using transcriptomic data generated from mouse placenta at e7.5, e8.5 and e9.5, we identified timepoint-specific gene groups that can be used for gene network inferences and analyses, as well as cell population annotations. Importantly, we were able to infer cell populations at different timepoints without known marker genes or reference dataset from the same species. The cell proportion inferences were necessary to bypass the confounding factors from cell heterogeneity, and thus predict more accurate novel regulators of cell-specific processes such as TB cell migration. This computational pipeline could be used to infer and analyze gene networks governing the development of placenta at other timepoints or to study developmental processes in other tissues.

We carried out DEA across the three timepoints on both the transcript and the gene level. These analyses revealed that a gene may have transcripts that are differentially expressed at different timepoints. For example, Igf2, a placental nutrient transport marker [76], has different transcripts grouped to e8.5 and e9.5 (Supplementary Table S1). This observation also aligned with a recent study which showed in 6–10 weeks’ and 11–23 weeks’ human placenta, differentially expressed genes, transcripts or differential transcript usage could all assist in the understanding of placental development [77]. Therefore, in future studies, investigating roles of both genes and their transcripts could give a more complete functional profile at each timepoint. Moreover, our results, together with previous studies in human placenta [23], [24], [77], suggest that time series transcriptomic analyses could be a useful approach to identify genes governing the development of the placenta. It will be beneficial to integrate these time series datasets to determine species-specific biomarkers of placental development.

We identified hub genes and their immediate neighboring genes which could regulate placental development and confirmed the regulatory roles of four novel genes (Mtdh, Siah2, Hnrnpk and Ncor2) in cell migration in the HTR-8/SVneo cell line. Interestingly, all four genes have been shown to have roles in cancer cells: Siah2 was shown to promote cell invasiveness in human gastric cancer cells by interacting with ETS2 and TWIST1 [78]; Mtdh regulates proliferation and migration of esophageal squamous cell carcinoma cells [79]; absence of Hnrnpk reduces cell proliferation, migration and invasion ability in human gastric cancer cells [80]; and repression of NCOR2 and ZBTB7A increased cell migration in lung adenocarcinoma cells [81]. This result further supports previous studies that show the comparability between placental cell migration and invasion, and tumor cell migration and invasion [71], [82], although specific genes may have different impacts on migration/invasion capacity such as with the Ncor2 gene. We acknowledge the HTR-8/SVneo cell line bears certain differences to TB cells such as in their miRNA expression profiles [83]. Therefore, in order to determine the exact roles of these genes in the placenta, future experiments such as migration assays in human TB stem cells derived with the Okae protocol [75] or gene knock out experiments in vivo would be necessary.

In our analyses, we observed that timepoint-specific genes and their networks represented expression profiles for specific placental cell populations at the three timepoints. In particular, analysis of e7.5-specific and e8.5-specific genes and networks showed that placental tissues at e7.5 and e8.5 contain different populations of TB cells, while e9.5-specific genes and networks showed multiple cell types including TB, endothelial and fibroblast cells. The significant overlap between e7.5-specific genes and genes of EVT cells yielded an interesting suggestion that the TB cell populations in e7.5 mouse placenta may share similarity in gene profiles to human EVT, although mouse TB and human EVT have certain differences such as their invasiveness levels [9]. Examples of EVT genes present in e7.5-specific gene group include FSTL3 (downregulation decreased TB migration and invasion in JAR cell line [84]), ADM (increased TB migration and invasion in JAR and HTR-8/SVneo cell line [85]), and ASCL2 (regulates TB differentiation [6]).

Furthermore, while it is true that single-cell (sc) resolution data is necessary to gain more information about the cell populations in the tissues, these results showed strong evidence that bulk RNA-seq data could be used to infer the cell type composition. In addition, scRNA-seq assays could be noisier than bulk RNA-seq due to various technical aspects such as the amount of starting materials, cell size, cell cycle, and batch effects [86], [87], which are difficult to be corrected for [88]. Therefore, bulk RNA-seq, ideally in conjunction with scRNA-seq, is beneficial for the study of biological processes that involve multiple cell types.

In our network analysis, we observed that the GO term “inflammatory response” was enriched in e7.5_1_STRING (q-value = 1.52E-23), e7.5_2_GENIE3 (q-value = 0.00012) and e9.5_2_STRING (q-value = 4.17E-10) (Supplementary Table S6). The inflammatory process could be happening in the placenta during e7.5 to e9.5 when TB cells actively invade the decidua [89] and create a pro-inflammatory environment [90]. Another possibility is contamination from decidual cells, which could be detected when combining bulk and scRNA-seq [91]. This further demonstrates the benefits of bulk and scRNA-seq data integration.

Upon conclusion of this study, we have shown that in the mouse placenta at e7.5, e8.5 and e9.5, genes with timepoint-specific expression patterns can be associated with distinct processes and cell types. The genes identified by timepoint-specific gene-network analysis could be interesting candidates for future studies focused on the understanding of placental development and placenta associated pregnancy disorders.

## Materials and Methods

### 1. RNA-seq library preparation and sequencing

Placenta tissue was collected from timed-pregnant CD-1 mice (Charles Rivers Labs) following the guidelines and protocol approved by Iowa State University Institutional Animal Care and Use Committee (IACUC), protocol number 18–350. Placenta samples were collected as previous described [19], [104] at e7.5, e8.5, and e9.5 and the age of the embryo was determined by following the embryonic development guidelines [105]. Briefly, tissues from the ectoplacental cone (EPC) and chorion were separated from the decidua, yolk sac, umbilical cord, and embryo, and then collected. For e7.5, 12 EPCs were collected and pooled into one replicate, as described in [21]. For e8.5, five placentas were collected per replicate, and for e9.5, one placenta was collected per replicate. Each timepoint had a total of 6 biological replicates.

Tissues were processed for RNA isolation immediately after collection using the Purelink RNA micro scale kit (Thermofisher, 12183016). RNA concentration and RIN values were measured using the RNA 6000 Nano assay kit on the Agilent 2100 Bioanalyzer (GTF facility, ISU), and all samples had a RIN score ≥ 7.7 (Supplementary Table S12). Further processing of the samples, library preparation and sequencing was performed by the DNA facility at Iowa State University. Libraries were sequenced using the Illumina HiSeq 3000 with single-end 50 base pair reads. The pooled library sample was run over two sequencing lanes (technical replicates for each sample).

### 2. RNA-seq data processing

The quality and adapter content were assessed using FastQC (version 0.11.7) [106]. Low quality reads and adapters were trimmed with Trimmomatic (version 0.39) [107].

Technical replicates were then merged, and the reads were pseudo-aligned and quantified (in TPM) using Kallisto (version 0.43.1; l = 200, s = 30; b = 100) [108]. Transcript sequences on autosomal and sex chromosomes of the mouse genome (GRCm38.p6) from Ensembl release 98 [109] were used to build the Kallisto index.

For further quality control, we carried out hierarchical clustering and principal component analysis (PCA) of samples. First, from the transcripts with raw counts ≥ 20 in ≥ 6 samples, we obtained the top 50% most variable transcripts, then centered and scaled their expression. Next, we implemented hierarchical clustering with the hclust() function in R (package stats [110], version 3.6.3), using the agglomerative approach with Euclidean distance and complete linkage. To implement PCA, we used the prcomp() function in R (package stats, version 3.6.3). We observed samples of each timepoint cluster close to each other and away from other timepoints. Outlier samples, which did not cluster with their respective timepoint groups, were removed prior to carrying out downstream analyses (Supplementary Figure S6).

### 3. Cluster analysis

Before performing all clustering procedures, transcripts with low raw counts (mean raw counts < 20 in all timepoints) were filtered out, and expression data (in TPM) was scaled and re-centered. Hierarchical clustering, k-means clustering, self-organizing map and spectral clustering was performed on the top 75% most variable protein coding transcripts (23,571 transcripts total).

To do hierarchical clustering, we implemented hierarchical clustering with the hclust() function in R (package stats [110], version 3.6.3), using the agglomerative approach with Euclidean distance and complete linkage. The resulting dendrogram was cut at the second highest level to obtain three clusters.

K-means clustering was carried out using the R function kmeans() (centers = 3, other parameters: default; package stats, version 3.6.3).

Self-organizing map clustering was performed with the R function som() with rectangular 3 × 1 grid (other parameters: default; package kohonen [111], version 3.0.10).

To implement spectral clustering, we utilized the following functions in R: computeGaussianSimilarity() (sigma = 1) to compute similarity matrix, and spectralClustering() (K = 3, other parameters: default; package RclusTool [112], version 0.91.3) to cluster.

To determine how the genes in each cluster relate to specific processes of placental development, we obtained gene lists from previously published review articles [5], [15]–[18], then calculate the percentage of markers in hierarchical clusters as (number of markers in a cluster)/(total number of markers of the process) × 100.

### 4. Differential expression analysis (DEA)

DEA at transcript and gene levels were carried out with Sleuth (version 0.30.0) [113] using the likelihood ratio test (default basic filtering) and the p-value aggregation process [114]. Fold change of a transcript was calculated using its average raw TPM across all samples. A transcript was considered differentially expressed (DE) if it had a fold change ≥ 1.5 and a q-value ≤ 0.05. A gene was considered DE if its q-value was ≤ 0.05 and had at least one protein-coding DE transcript. For lists of DE protein-coding transcripts that had at least one DE gene, and lists of DE genes with at least one DE protein-coding transcripts, see Supplementary Table S3.

### 5. Definition of timepoint-specific genes

Timepoint-specific gene groups are defined as the following:

1. E8.5-specific transcripts: transcripts in e8.5 hierarchical cluster, are up-regulated at e8.5 (compared to e7.5) or are up-regulated at e8.5 (compared to e9.5). E8.5-specific genes are ones associated with e8.5-specific transcripts.
2. E7.5-specific transcripts: transcripts in e7.5 hierarchical cluster, are up-regulated at e7.5 (compared to e9.5), and are not in e8.5-specific group. E7.5-specific genes are ones associated with e7.5-specific transcripts.
3. E9.5-specific transcripts: transcripts in e9.5 hierarchical cluster, are up-regulated at e9.5 (compared to e7.5), and are not in e8.5-specific group. E9.5-specific genes are ones associated with e9.5-specific transcripts.

### 6. Network construction and analysis

The STRING database (version 11.0b) [35] was used to build protein – protein interaction networks at each timepoint. Edges from evidence channels: experiments, databases, text-mining and co-expression with confidence score ≥ 0.55 were chosen for further analyses.

Gene regulatory networks at each timepoint were constructed with GENIE3 (version 1.16.0) [36]. At each timepoint, as inputs for GENIE3, timepoint-specific transcripts with average TPM at the timepoint ≥ 5 were aggregated to obtain gene counts with the R package tximport (version 1.14.2; countsFromAbundance = lengthScaledTPM) [115]. Genes that encode transcription factors (TFs) and co-TFs, downloaded from AnimalTFDB (version 3.0) [116], were treated as candidate regulators. Then, edges with weight < the 90^th^ percentile were filtered out.

Largest connected components of the networks were analyzed using Cytoscape (version 3.7.2) [117]. All networks were treated as undirected, and network sub-clustering was performed using the GLay plug-in (default parameters) [37]. Networks with ≥ 100 nodes were used for further analyses. Hub genes were defined as nodes that have degree, betweenness and closeness centralities in the 10^th^ percentile of their networks.

To determine the relevant functions of the genes in networks, we used gene ontology (GO) analysis. ClusterProfiler (version 4.0.5) [118] was used, with the mouse annotation from the org.Mm.eg.db R package (version 3.13.0) [119], the maximum size of genes = 1000, and a q-value cut-off = 0.05. Next, a fold change for each term was calculated as GeneRatio/BgRatio. A GO term was considered enriched when its q-value ≤ 0.05, fold change ≥ 2, and the number of observed genes ≥ 5.

A gene was determined to have an annotated role in placental development if it was annotated with experimental evidence or had an associated PubMed ID under all GO and MGI Phenotype terms related to placenta, TB cells, TE and chorion layer. Experimental evidence codes include “Inferred from Experiment” (EXP), “Inferred from Direct Assay” (IDA), “Inferred from Physical Interaction” (IPI), “Inferred from Mutant Phenotype” (IMP), “Inferred from Genetic Interaction” (IGI), and “Inferred from Expression Pattern” (IEP). GO terms, MGI Phenotype terms and gene annotations were downloaded from MGI (http://www.informatics.jax.org/) (version 6.19) [38]. For lists of terms used, see Supplementary Table S7.

### 7. Deconvolution analysis

To infer the proportion of cell types across timepoints, we carried out deconvolution analysis using the R package LinSeed (version 0.99.2) [68]. Gene abundances (in TPM) used as inputs for the analysis were obtained using tximport (version 1.14.2; countsFromAbundance = lengthScaledTPM) [115]. Then, we used top 5000 most expressed genes across timepoints, and sampled 100000 times to test for the significance of the genes to be used for deconvolution analysis. A significant gene was one with p-value ≤ 0.05. The number of cell groups was determined after examining the singular value decomposition (SVD) plot, generated with the svdPlot() function in LinSeed. Cell markers were defined as the top 100 genes closest to the cell group’s corner, and closer to the corner than any other corners.

### 8. Placenta Cell Enrichment and Placenta Ontology analysis

The PlacentaCellEnrich webtool [69] and Placenta Ontology [71] were used to infer the relevant cell types using gene lists. For PlacentaCellEnrich, cell-type specific groups were based on the single-cell transcriptome data of human maternal-fetal interface from Vento-Tormo et al. [26]. A enrichment was considered significant if its adj. p-value is ≤ 0.05, fold change ≥ 2, and the number of associated genes found is ≥ 5. For Placenta Ontology, we obtained placenta ontology GMT file from Naismith et al. and uploaded the file to the WEB-based GEne SeT AnaLysis Toolkit (www.webgestalt.org) [120] as a functional database. An ontology with FDR ≤ 0.05, fold change ≥ 2 and the number of observed genes ≥ 5 was considered enriched.

### 9. *In vitro* validation experiments

#### Cell culture

HTR-8/SVneo (ATCC CRL3271) were cultured as recommended by ATCC and as done by others [121]. Briefly, cells were grown in RPMI-1640 media (ATCC 302001) supplemented with 5% FBS (VWR, 97068-085) without antibiotics. Cells were split every 3 to 4 days, at 80-90% confluency.

#### siRNA knockdown

HTR-8/SVneo cells were transfected with two different siRNA for each target gene knockdown (KD). Cells were split at 80% confluency, and siRNA transfection was performed in 6-well plates; 150,000 cells/well were seeded [20]. After 24 hours, cells were transfected with 30nM siRNA using RNAiMax 3000 (Thermofisher, 13778150). Media was replaced after 24 hours of transfection, and cells were collected after 48 hours of transfection and seeded for migration assays. GAPDH was used for normalization of all four genes’ expression (ΔCT). Percent KD were calculated with the ΔΔCT method. SiRNA and primer information can be found in Supplementary Table S12.

#### Migration assays

Migration assays were performed using Costar inserts (Corning, 3464). The inserts were placed in a 24 well plate and 75,000 cells in serum-free RPMI media (ATCC, 30-2001) were directly seeded in the top chamber of the insert. The bottom chamber was filled with 600ul of RPMI media supplemented with 10% FBS as a chemoattractant. The cells were allowed to migrate for 24 hours at 37°C. The cells on the bottom of the inserts were fixed in 4% PFA (Fisher Scientific, AAJ61899AK) for 5 min and then washed for 1 min with PBS twice. The cells in the top chamber were scraped off using a wet q-tip (Fisher Scientific, 22029488) and the cells on the bottom of the inserts were stained with Hematoxylin (Fisher Scientific, 23245677) for 24 hours. The inserts were washed twice in distilled water. The membrane was cut using a scalpel (Fisher Scientific, 1484002) and mounted on a clean glass slide in Vectamount mounting medium (Fisher Scientific, NC9354983). The cells were observed under a dissection microscope and imaged at 12.5X magnification. The images were analyzed using the ImageJ tool, and the integrated density was obtained for each image.

#### Statistical analysis

Experiments were performed with three replicates per condition (negative control or knockdown) per gene. P-values were calculated with Wilcoxon rank sum test.

## Data Availability Statement

All code for the analyses is available at https://github.com/Tuteja-Lab/PlacentaRNA-seq. All raw and processed data is available for download on NCBI Gene Expression Omnibus (GEO) Repository, accession number: GSE202243.

Acknowledgements

We acknowledge the Iowa State University DNA Facility for preparing the library and providing sequencing for RNA-seq experiments, and the Research IT group at Iowa State University (http://researchit.las.iastate.edu) for providing servers and IT support. We would like to thank Tuteja lab members for their discussion and support.

## Author contributions

Conceptualization, H.V., G.T.; Methodology, H.V., G.T.; Data Generation, H.K., R.S., Formal Analysis, H.V.; Data Interpretation, H.V., G.T.; Experimental Validation, H.K.; Analysis Validation, K.K.; Writing – Original Draft Preparation, H.V., G.T.; Writing – Review and Editing, H.V., H.K., K.K., R.S. and G.T.; Supervision, G.T.; Funding Acquisition, G.T.

## Abbreviations

e: Embryonic day
RNA-seq: RNA sequencing
TB: Trophoblast
TE: Trophectoderm
EPC: Ectoplacental cone
TGC: Trophoblast giant cells
GO: Gene ontology
MGI: Mouse Genome Informatics
TPM: Transcripts per million
EVT: Extravillous trophoblast
SCT: Syncytiotrophoblast
VCT: Villous trophoblast
adj. p-value: Adjusted p-value
FDR: False discovery rate
sc: Single-cell
DEA: Differential expression analysis
DE: Differentially expressed
FPKM: Fragments per kilobase of transcript per million of mapped reads

