## Supplementary Figures for "Identifying novel regulators of placental development using time series transcriptomic data and network analyses"

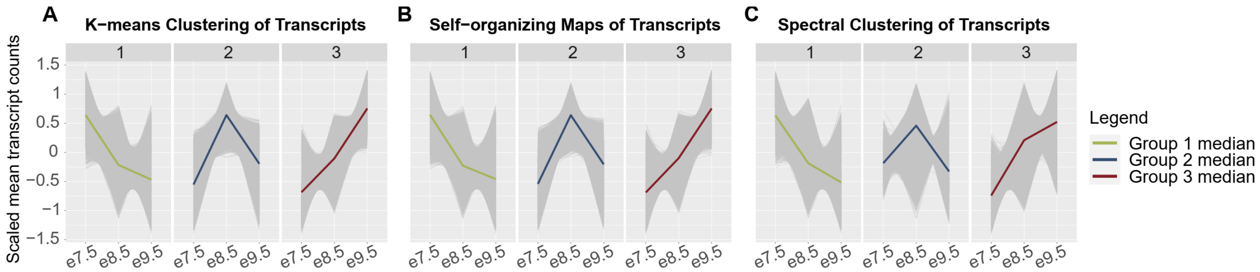


Figure S1: Clustering of transcripts using different methods

Line charts of scaled mean raw counts of transcripts in (A) K-means clusters, (B) self-organizing maps, and (C) spectral clusters, showing group median expression levels peak at each timepoint, agreeing with results from hierarchical clustering (Figure 1B). Green: median expression of transcripts in group 1; blue: median expression of transcripts in group 2; dark red: median expression of transcripts in group 3.

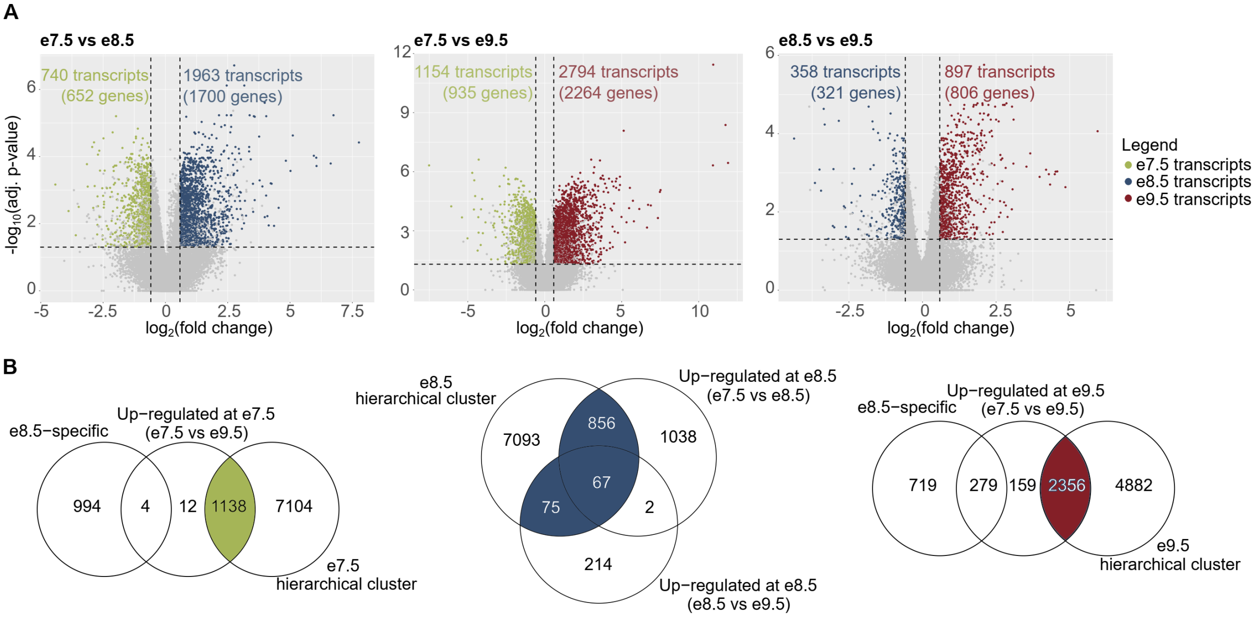


Figure S2: Identification of timepoint-specific genes using the combined results from hierarchical clustering and differential expression analyses

A. Volcano plots showing the number of differentially expressed (DE) transcripts in each analysis: e7.5 vs e8.5, e8.5 vs e9.5, e7.5 vs e9.5. A transcript is DE if its q-value ≤ 0.05 and fold change ≥ 1.5. Green: transcripts up-regulated at e7.5; blue: transcripts up-regulated at e8.5; dark red: transcripts up-regulated at e9.5.
B. Venn diagrams showing definitions of timepoint-specific transcripts (see Methods). There are 1138 e7.5-specific transcripts (922 genes), 998 e8.5-specific transcripts (915 genes), and 2156 e9.5-specific transcripts (1952 genes). Green: e7.5-specific transcripts; blue: e8.5-transcripts; dark red: e9.5-specific transcripts.

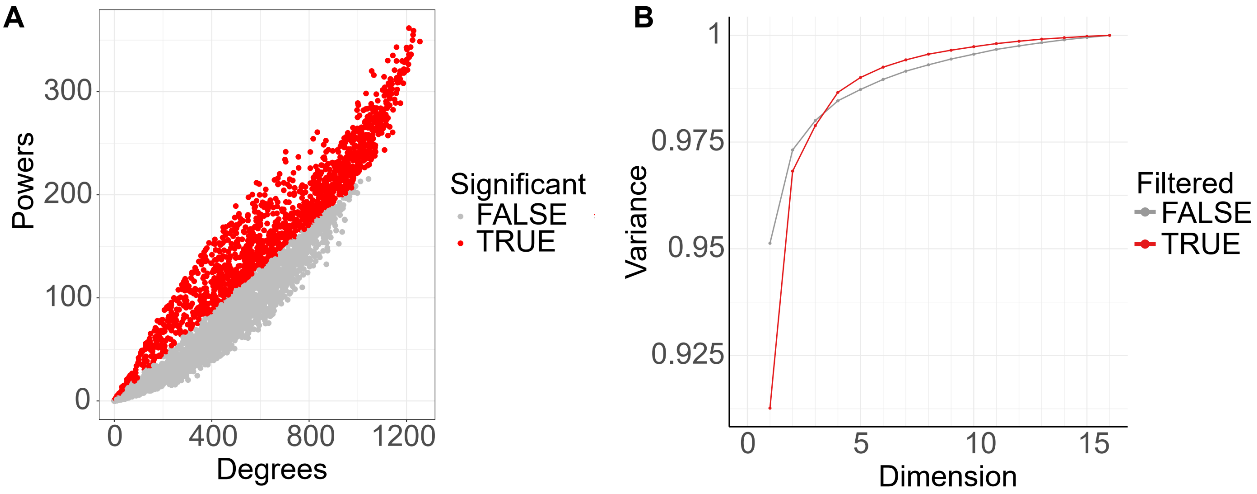


Figure S3: Deconvolution analysis results

A. 1413 genes (colored red) were determined to be significant for deconvolution analysis. Starting with top 5000 most expressed genes across timepoints (expression in TPM), we used LinSeed to sample 100000 times in order to determine the significant genes (p-value ≤ 0.05) for the gene linearity network topology.
B. Singular value decomposition plot showing the amount of variance the significant genes (red line) could capture with different numbers of dimensions (number of cell groups). Five cell groups, which captured 99% of the variance, were determined as present the samples.

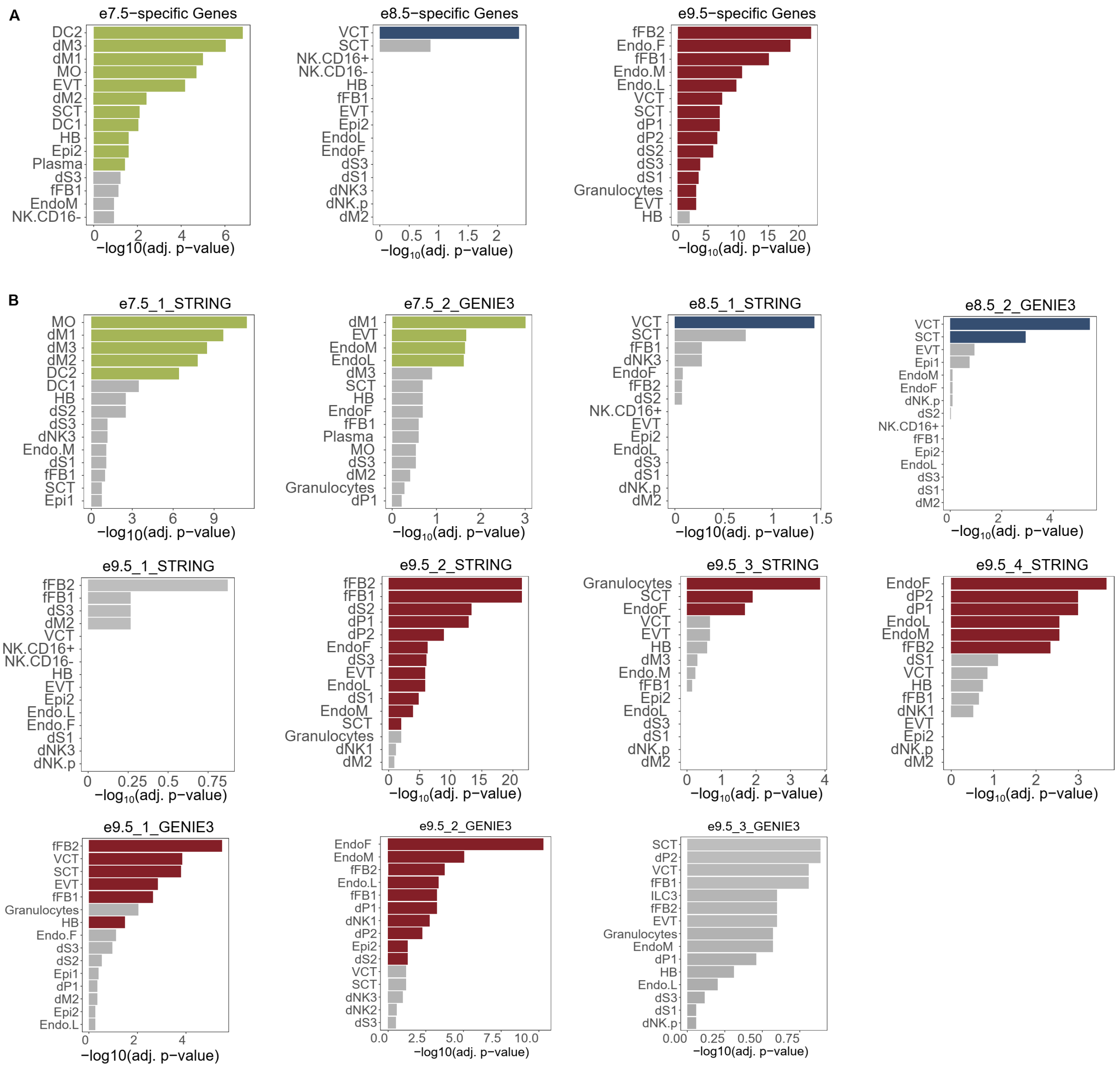


C.

Figure S4: Full PlacentaCellEnrich (PCE) results.

Colored bars: cell types with significant enrichments; grey bars: cell types with insignificant enrichments. The enrichment was significant if adj. p-value ≤ 0.05, fold change ≥ 2 and number of observed genes ≥ 5.

A. Bar plots showing the cell-type specific enrichments of mouse timepoint-specific gene sets.

B. Bar plots showing the cell-type specific enrichments of subnetwork genes.

Abbreviations: SCT, syncytiotrophoblast; HB, Hofbauer cells; EVT, extravillous trophoblast; VCT, villous cytotrophoblast; EndoF, fetal endothelium; fFB1, fetal fibroblast cluster 1; fFB2, fetal fibroblast cluster 2; DC, dendritic cells; dM, decidual macrophages; dS, decidual stromal cells; EndoL, lymphatic endothelium; Epi, epithelial glandular cells; NK, natural killer cells.

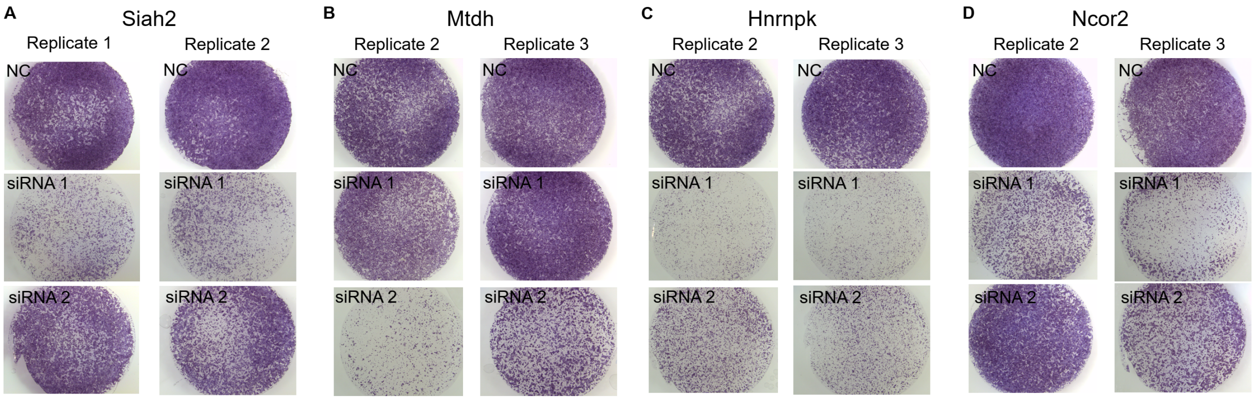


Figure S5: Images of cell migration assays for all replicates not shown in Figure 4.

A. *Siah2* experiments. B. *Mtdh* experiments. C. *Hnrnpk* experiments. D. *Ncor2* experiments.

**A**

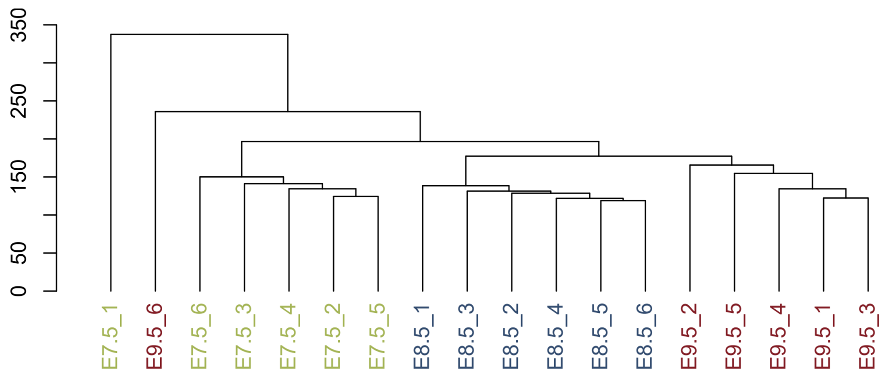


**B**

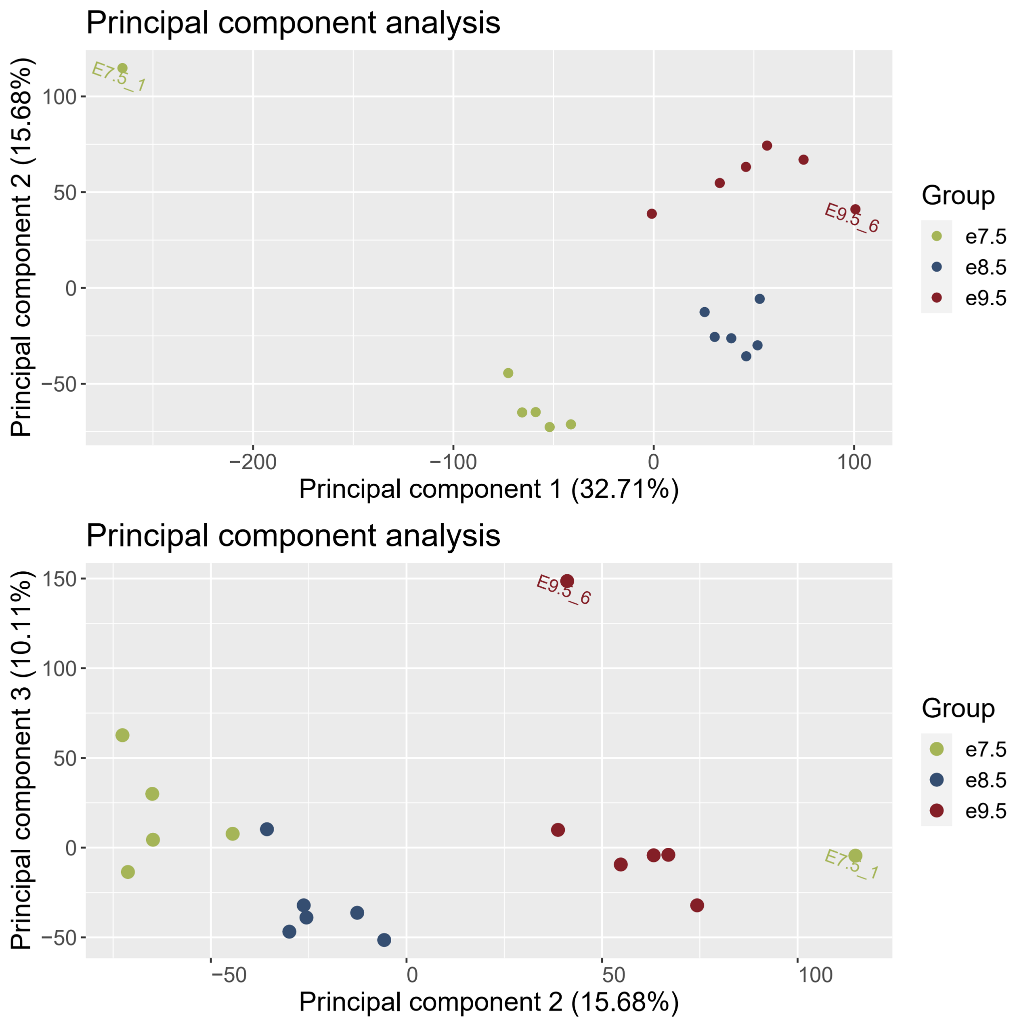


Figure S6: Quality control of samples. Samples e7.5_1 and e9.5_6 were removed prior to downstream analysis due to their large distance from their timepoint's group (see Methods).

A. Hierarchical clustering of samples.

B. Principal component (PC) analysis of samples. Panel 1: PC2 vs PC1; panel 2: PC3 vs PC2.
